# A generative language model decodes contextual constraints on codon choice for mRNA design

**DOI:** 10.1101/2025.05.13.653614

**Authors:** Marjan Faizi, Helen Sakharova, Helene Borrmann, Mohammad S. Parsa, Liana F. Lareau

## Abstract

Synonymous codon choice affects mRNA fate and protein output, posing a challenge for mRNA technology. Design of therapeutic mRNAs requires a model that captures biological nuance from natural sequences while identifying constraints relevant to synthetic expression. We present Trias, a generative encoder-decoder language model trained on millions of vertebrate coding sequences that successfully designs mRNAs with high protein output. Trias learns codon usage rules directly from sequence data, integrating local and global dependencies to generate species-specific codon sequences that reflect biological constraints. Without explicit training on protein expression, its zero-shot predictions correlate strongly with experimental measurements of mRNA stability and protein output. Moreover, we show experimentally that novel sequences generated by Trias for a therapeutically relevant target produce more protein than existing methods, demonstrating that preferences learned from natural sequences can lead to high output from synthetic sequences. Released open source with full code and weights, Trias provides a reproducible framework for synthetic mRNA design that reveals molecular and evolutionary principles behind codon choice.

## Introduction

Synthetic messenger RNA technology has emerged as a transformative tool for biotechnology, opening new frontiers in synthetic biology, genome engineering, and medicine. mRNAs encoding proteins of interest can be delivered into cells for translation by the cell’s own machinery. A key challenge is to design mRNA sequences that drive high levels of heterologous protein expression. Natural mRNAs are shaped by overlapping functional constraints spanning the transcript from the 5’ cap through the untranslated regions, coding sequence, and polyA tail, all subject to optimization for synthetic mRNA design (Sample *et al*., 2019; Lewis *et al*., 2025; Leppek *et al*., 2022; Zhang *et al*., 2023).

Design of the coding sequence presents a particular challenge. While the desired protein sequence may be pre-determined, the choice between synonymous codons for each amino acid becomes a combinatorially complex problem. For example, the SARS-CoV-2 spike protein can be encoded by more than 10^632^ different codon sequences (Kim *et al*., 2024). Within this vast range of possibilities, codon choice is not random; organisms favor certain codons, and these preferences vary by species. Yet, the underlying rules guiding optimal codon selection remain poorly understood (Wu and Bazzini, 2023).

Codon choice has subtle but powerful impacts on the outcomes of translation. Common codons are decoded faster and more accurately due to more abundant tRNAs, while rare codons slow translation (Liu, 2020). Slow translation can destabilize mRNAs through codon optimality mediated decay (Presnyak *et al*., 2015; Bazzini *et al*., 2016; Narula *et al*., 2019). It can also reduce the number of proteins made per mRNA; codon choice alone can create an eight-fold range in protein output (Tunney *et al*., 2018; Barrington *et al*., 2023; Lyons *et al*., 2026). However, fast translation is not always ideal. Multi-domain proteins can require precisely timed translation for proper folding and function (Kim *et al*., 2015; Yu *et al*., 2015; Kimchi-Sarfaty *et al*., 2007; Kirchner *et al*., 2017).

Codon choice also shapes mRNA properties beyond the process of translation. The higher GC content of preferred codons in mammalian genes leads to more stable mRNA structure, affecting many aspects of an mRNA’s fate. GC content also influences mRNA processing and nuclear export (Zuckerman *et al*., 2020; Mordstein *et al*., 2020). Coding sequences are further overlaid with splicing motifs and other regulatory elements. Many of these effects depend on nuclear processes that would not be encountered by a synthetic mRNA delivered to the cytoplasm, but synthetic mRNAs can have distinct constraints. Higher GC content promotes greater stability in solution even before an mRNA vaccine is administered, affecting how much intact mRNA survives to reach the ribosome (Leppek *et al*., 2022).

These observations underscore the need for sophisticated approaches to sequence design that consider biological context. Most commercial tools rely on manually defined features and cannot fully capture the dependencies between codons. Deep learning approaches that learn directly from experimental data on ribosome distribution represent a major step forward, but they may not capture complex sequence dependencies (Tunney *et al*., 2018; Jain *et al*., 2023). Large language models represent a major advance in machine learning, and the same powerful approach can be applied directly to the language of biological sequences. Sequence language models of protein or DNA learn the underlying distributions and patterns from millions of naturally occurring sequences (Simon *et al*., 2024). They can serve as foundation models that generalize across a wide range of biological tasks including structure prediction and variant effect prediction, either in a zero-shot setting (without fine-tuning or task-specific examples) or after fine-tuning (Rives *et al*., 2021; Lin *et al*., 2023; Benegas *et al*., 2023; Karollus *et al*., 2024; Nguyen *et al*., 2024a,b; Brixi *et al*., 2026). Building on the success of these models, specialized codon language models have been trained on comprehensive datasets of mRNA coding sequences (Li *et al*., 2024; Constant *et al*., 2023; Outeiral and Deane, 2024; Sidi *et al*., 2024; Fallahpour *et al*., 2025; Naghipourfar *et al*., 2024; Zhang *et al*., 2025). These broad models have the potential to uncover complex codon usage rules that traditional methods may not capture, providing an improved tool for codon optimization and a framework for studying synonymous codon choice.

Recent codon models have progressed from assessing the properties of existing sequences toward designing new sequences with generative architectures (Fallahpour *et al*., 2025; Constant *et al*., 2023; Sidi *et al*., 2024; Naghipourfar *et al*., 2024). Here we present a new model, Trias, that employs a powerful encoder-decoder architecture well suited for sequence generation. It expands the capability of previous models by enabling design of full-length sequences of up to 2,046 amino acids and by focusing on vertebrate rather than microbial sequences, capturing their distinct sequence constraints to enable design for expression in human cells. In parallel with our present work, GEMORNA (Zhang *et al*., 2025) introduced a conceptually similar encoder–decoder codon model trained on mammalian coding sequences with long-context modeling capabilities, reflecting similar advances in generative modeling of codon usage.

Our work validates the implicit assumption that a model trained on natural sequence can design sequences for high output for vaccines and therapeutics. Powerful language models like Trias hold great promise for synthetic sequence design, but only if they align with the optimization tasks to which they are applied. Foundation models capture the distribution of natural sequences that have been shaped by evolution for proper production, processing and function but not necessarily for maximal expression. Understanding what these models learn about natural codon usage and how this knowledge translates to synthetic applications is critical for their successful use in sequence design.

We show that Trias learns to design sequences with better-than-natural codon usage while recognizing some proteins that employ less-preferred codons. With no training on experimental data, zero-shot predictions from Trias correlate strongly with experimental measurements of protein expression and mRNA stability across independent studies spanning distinct reporter genes in human cells and in mouse. Trias outperforms competing models at these zero-shot predictions. Further, Trias generates new sequences encoding therapeutically relevant proteins that produce higher output than existing methods. Trias provides a data-driven method for codon optimization in recombinant protein production and mRNA-based therapeutics as well as a tool for studying how codon usage is shaped by sequence context. To facilitate its broad application, we have released Trias on Hugging Face, where it enables reverse translation of protein sequences into codon sequences for over 600 vertebrate species.

## Results

### Trias learns codon preferences and amino acid properties through generative training

Trias is a sequence-to-sequence model designed to reverse translate protein sequences into codon sequences (Figure 1A). It is based on the BART architecture with bidirectional encoder and autoregressive decoder modules, commonly used in natural language processing tasks such as machine translation (Lewis *et al*., 2019). The bidirectional encoder module of Trias processes the entire input protein sequence at once, capturing long-range dependencies between amino acids. It produces a latent representation of the sequence that is passed to the decoder. The autoregressive decoder then generates the corresponding codon sequence token by token, conditioning each predicted codon on the previously generated codons and the encoder’s representation. The model is trained by providing the encoder module with a corrupted version of each protein sequence with 15% of amino acid positions masked and then calculating the reconstruction loss of the generated codon sequence against the true codon sequence. To allow the model to learn species-specific preferences for codon usage, we prepend species tags to the protein input sequence.

**Figure 1:**
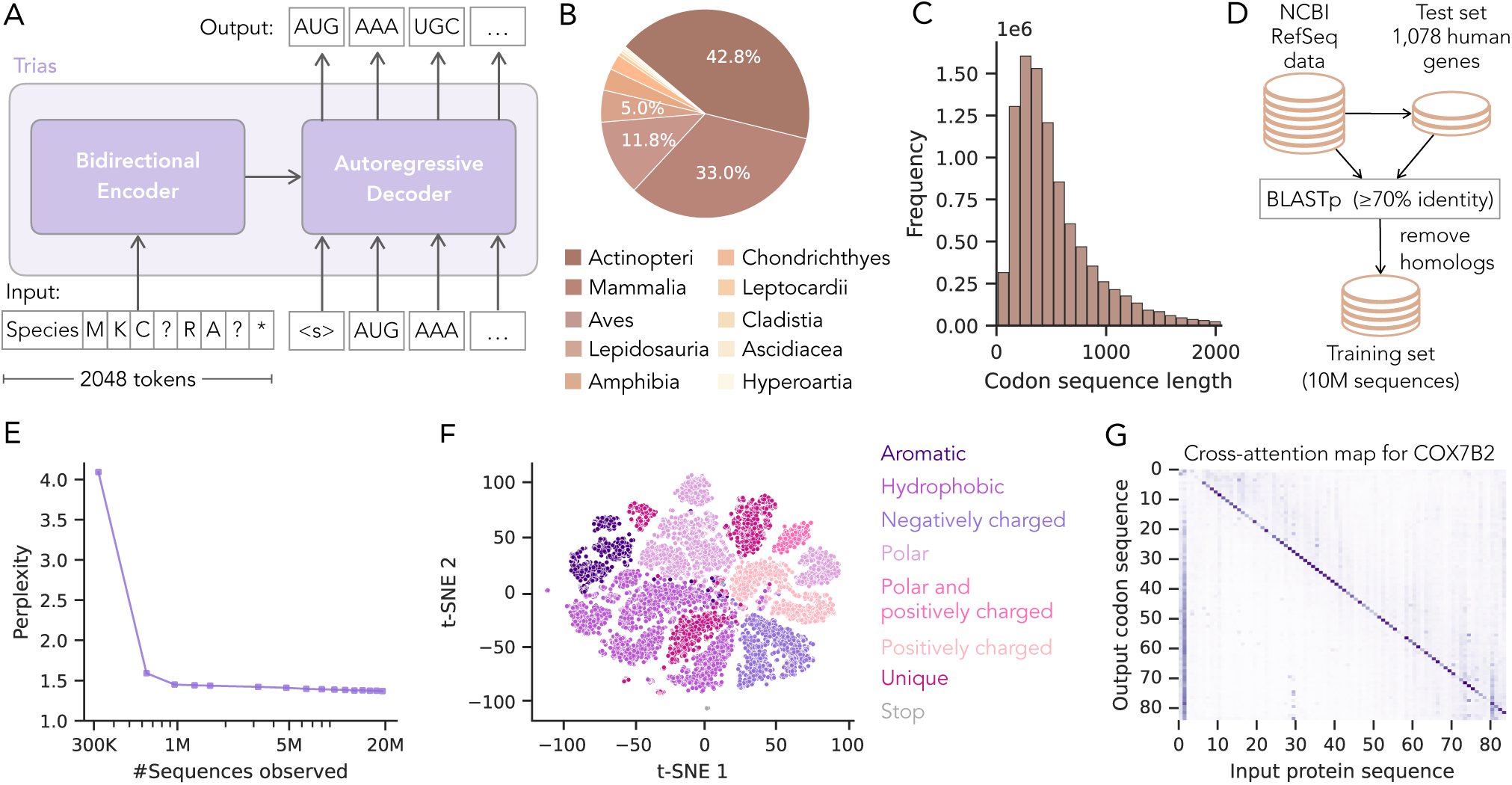
Model overview, dataset composition and training performance. (A) Trias is an encoder-decoder model that reverse translates protein sequences into codon sequences while incorporating species-specific codon preferences. During training, 15% of input tokens are randomly masked (indicated as ’?’). (B) The model was trained on 10M codon sequences from 640 vertebrate species. (C) Distribution of codon sequence lengths in the dataset. (D) A test set of 1,078 human genes was set aside and their close mammalian homologs (*≥* 70% protein identity) were removed from the training set. (E) Perplexity during training, showing a decrease from 4.1 to 1.4 on the training set. (F) Projection of the final encoder embedding layer for 20 randomly selected human protein sequences, with each dot representing an amino acid. (G) Cross-attention map for example gene, showing how each codon attends to the input amino acid sequence and the species tag. Each row corresponds to a codon, and each column to an amino acid. The example shown represents a single attention head, averaged across all layers.

Trias has 47 million parameters and was trained on 10 million sequences from 640 vertebrate species sourced from the NCBI RefSeq database (Figure 1B), with sequences up to 2,046 tokens long (Figure 1C). Although language models do not typically employ a held-out test set for evaluation, we hoped to compare Trias-generated and natural sequences directly, necessitating careful construction of a test set of protein sequences never seen by Trias. At evolutionary distances further than the root of the mammalian lineage, conserved amino acid positions are unlikely to have conserved codon choice in the absence of selection (Warren *et al*., 2008), while genes with a more recent evolutionary divergence might have conserved codon choice because mutations have not yet saturated. We therefore set aside a test set of 1078 human proteins and then removed their close mammalian homologs from the training set, leaving only homologs distant enough that codon choice is no longer expected to be conserved (Figure 1D). This choice was intended to allow the model to learn recurrent patterns, such as a propensity for certain codons in particular types of protein domain, without memorizing specific examples.

We evaluated training convergence by tracking perplexity, quantifying the average number of codon choices the model considers per amino acid to show the model’s uncertainty in predicting codons (Figure 1E). During the first 640K examples, perplexity dropped from 4.1 to 1.6, indicating that the model quickly learned synonymous codon preferences. We also confirmed that, although Trias was not explicitly provided with the codon table, it quickly learned to select valid synonymous codons for each amino acid (Supplementary Figure S1 and S2).

We confirmed that the encoder module learned meaningful latent representations of amino acids by visualizing the t-SNE projection of the final encoder hidden layer for 20 randomly selected human protein sequences (Figure 1F, Supplementary Figure S3). Amino acids clustered by biochemical properties, similar to ESM-1b (Rives *et al*., 2021), with distinct groups for hydrophobic, polar, and charged residues. Next, to illustrate how Trias conditions its codon predictions on protein context, we visualized cross-attention weights for an example gene, COX7B2 (Figure 1G, Supplementary Figure S4). Cross-attention indicates how much each predicted codon attends to different positions in the input protein sequence. The strong diagonal signal indicates that codon choice is primarily influenced by the identity of its target amino acid, as expected. The vertical band at the start highlights the species tag, which influences codon selection across the sequence. Off-diagonal signals show that the model also attends to distant residues, suggesting that Trias incorporates both local and long-range context during codon generation.

### Trias captures preferences beyond codon frequency

Simple codon optimization methods favor frequent codons, but natural sequences deviate from these preferences in ways that may reflect functional constraints. An ideal model for designing synthetic mRNAs would capture the nuances necessary for proper function while enabling higher-than-natural protein output. The resulting sequence may resemble its natural counterpart in some respects and depart from it in others. To evaluate the generative performance of Trias, we generated codon sequences for each protein in the human test set (1078 sequences) using a greedy decoding strategy that selects the most probable codon at each step based on the previously generated sequence. We then compared these generated sequences to their wild-type counterparts across three interrelated properties: relative synonymous codon usage (sRSCU), GC content at the third codon position, and minimum free energy (MFE) of mRNA structure (Figure 2A).

**Figure 2:**
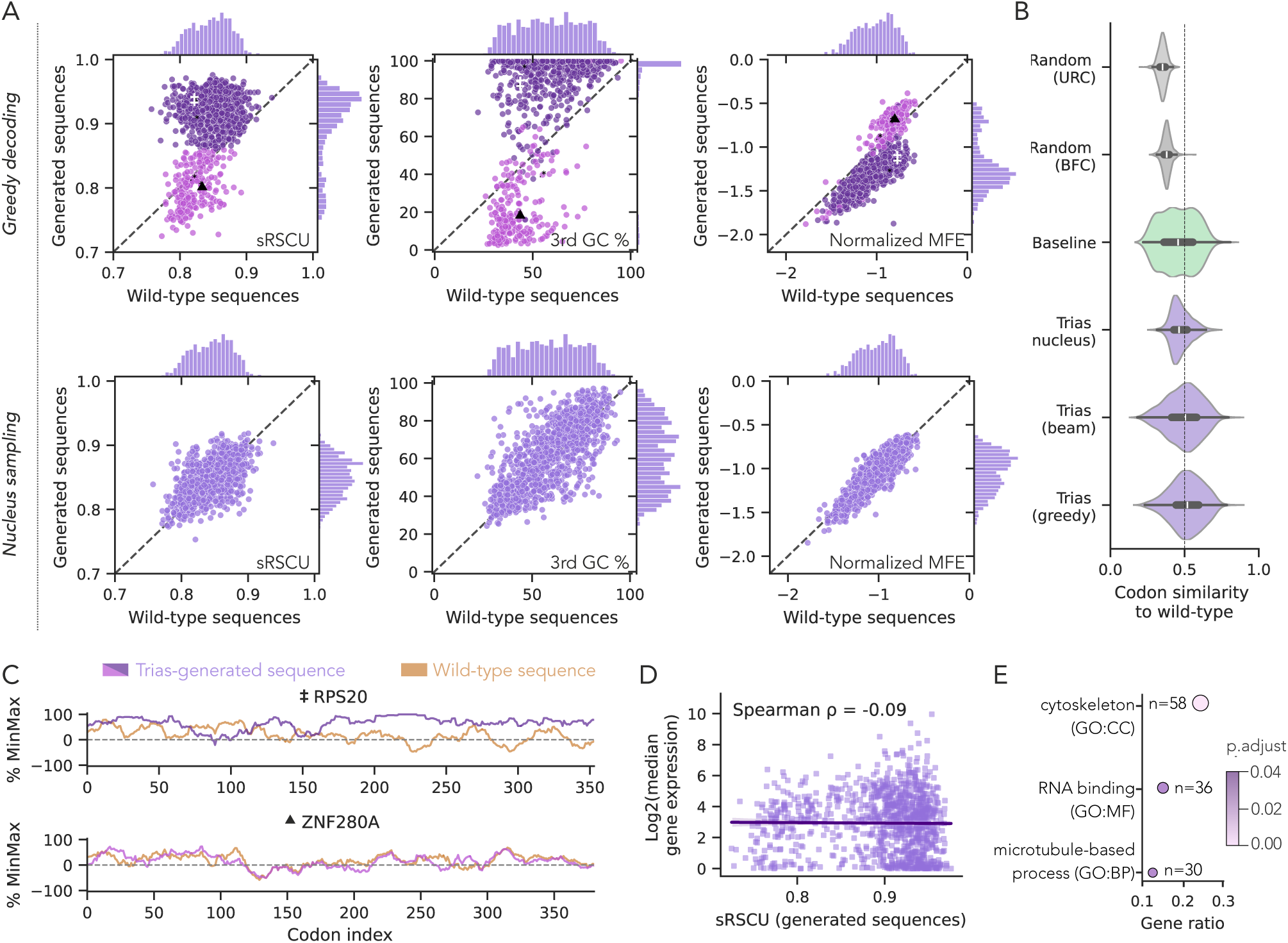
Sequence-level analysis of Trias-generated human codon sequences. (A) Comparison of Trias-generated sequences, using greedy decoding and nucleus sampling, to their wild-type counterparts across scaled relative synonymous codon usage (sRSCU), GC content at the third codon position, and minimum free energy (MFE, normalized by sequence length, computed using the ViennaRNA package (Lorenz *et al*., 2011)) in the human test set (1078 sequences). Sequences were colored purple or pink according to high or low generated sRSCU values to facilitate comparison across metrics. (B) Codon similarity between Trias-generated sequences and wild-type sequences, benchmarked against a baseline model selecting the most frequent codon at each position, uniform random synonymous codon selection, and random selection based on background codon frequencies. (C) %MinMax profiles illustrate codon usage fluctuations across genes with naturally low sRSCU, marked with symbols in panel A. ZNF280A maintains low sRSCU similar to wild-type, while RPS20 has higher predicted sRSCU. (D) Gene ontology (GO) enrichment analysis of genes with predicted low sRSCU (*≤*0.86). Shown are significantly enriched GO terms (adjusted p-value < 0.05) from the molecular function (MF), biological process (BP) and cellular component (CC) categories. (E) Log2-transformed median gene expression across all tissues in the GTEx dataset plotted against predicted sRSCU values for 1020 human test set genes found in GTEx.

Interestingly, although the naturally occurring sequences had a wide, unimodal distribution of sRSCU, the generated sequences fell into two populations (Figure 2A). The bulk of the sequences, spanning the full range of natural sRSCU values, were generated to have more preferred codons than their natural counterparts. This shift toward preferred codons may be desirable for synthetic sequences intended for high expression. A distinct, smaller set of sequences were generated with low sRSCU that matched the low natural sRSCU of these sequences. This outcome may be valuable for synthetic design if it reflects real patterns of constraint on these sequences. The third position GC content and MFE followed the same bimodal pattern as sRSCU, with sequences in the high-sRSCU population exhibiting higher GC content and stability.

The choice of decoding strategy shapes how closely the generated sequences resemble natural sequences. Greedy decoding selects the highest-probability codon at each step, which will tend to produce higher-than-natural sRSCU, as seen above. Beam search, which explores multiple high-probability continuations, yielded the same bimodal sRSCU and GC distributions (Supplementary Figure S5). Nucleus sampling, which selects probabilistically among likely codons, instead generated sequences whose sRSCU, GC content, and MFE distributions tracked the wild-type distributions with no distinct subpopulations (Figure 2A). The greedy approach may be more useful for designing optimized mRNAs for high expression, while nucleus sampling may suit applications that aim to preserve the diversity of natural sequences.

To test whether Trias captures sequence context beyond codon frequency, we compared its generative methods to three baseline mRNA design approaches: uniform random codon choice (URC), which samples uniformly from synonymous codons; background frequency codon choice (BFC), which samples according to species-specific codon usage frequencies; and a common codon baseline that always selects the most common codon for each amino acid. All three methods of generating sequences from Trias produced sequences that were more similar to their wild-type counterparts than those produced by the three baseline methods (Figure 2B; all pairwise comparisons *p <* 1 *×* 10*^−^*^5^, Mann-Whitney U test). As the baseline methods cannot capture any gene-or position-specific information, the improved performance of nucleus sampling vs the BFC baseline and of greedy decoding vs the common codon baseline shows that Trias learns a probability distribution that captures sequence context.

We then considered how the model size and training breadth affected predictions by comparing Trias to Trias-small, a model with fewer parameters (12.3 million) trained only on mammalian sequences and with the same heldout test set. Trias-small produced the same bimodal split in per-gene sRSCU, but more pronounced: sequences tended to be generated with uniformly optimal or uniformly rare codons, with fewer occupying the intermediate range (Supplementary Figure S6). Sequences generated by Trias-small also showed lower codon similarity to wild-type sequences (47%, compared to 52% for the full model). These differences suggest that the small model captures overall codon preferences but the broader training of the full model supports finer modulation of codon usage.

### Trias captures gene-level functional constraints on codon choice

We next asked what drives Trias to assign low sRSCU to some genes while optimizing others. Figure 2C shows two example genes with naturally low sRSCU treated differently by the model: ZNF280A, where Trias preserved rare codon usage, and RPS20, where it predicted more frequent codons than in the wild-type sequence. If a gene has functional constraints favoring non-preferred codons, this could appear as a conserved pattern of codon choice learnable from common protein sequence features across distantly related genes. The training and test set genes are distant enough that shared codon usage is not expected in the absence of functional constraint, avoiding simple memorization.

To see if Trias learned constraints for particular categories or families of genes, we performed a gene ontology (GO) analysis on the genes that Trias assigned low sRSCU (Figure 2D). These genes were enriched for RNA-binding function, microtubule-based processes, and cytoskeletal components. Genes with high predicted sRSCU showed no significant enrichment. A number of the low-sRSCU genes belonged to the large and highly conserved family of zinc finger proteins, suggesting that their codons might have constraints that are particularly learnable from the large number of distant examples.

A broader functional constraint could arise from pressure for high expression: in many model organisms, highly expressed genes tend to employ preferred codons, and expression alone can be a strong predictor of codon usage (Duret and Mouchiroud, 1999; Cope *et al*., 2018). Although Trias does not directly see information about expression, it could learn a preference for frequent codons, if present, across many different families of highly expressed genes that each have recognizable protein features. However, we found no significant correlation between predicted or natural sRSCU and mRNA expression levels from GTEx (Figure 2E, Supplementary Figure S7), in keeping with analyses showing that the relationship is weak or absent in mammals (Bénitière *et al*., 2025).

### Sequence context influences rare codon predictions

The previous analyses focused on the overall codon content per gene; we then considered factors influencing individual choices of rare or preferred codons within genes. Defining rare codons as those with sRSCU < 0.8 (Supplementary Figure S8), we classified all codon predictions from the human test set sequences into four categories: rare or common codons in the natural sequence that were predicted as rare or common codons by the model (Figure 3A). We then considered attributes of the sequence that potentially affected the outcome (Figure 3B).

**Figure 3:**
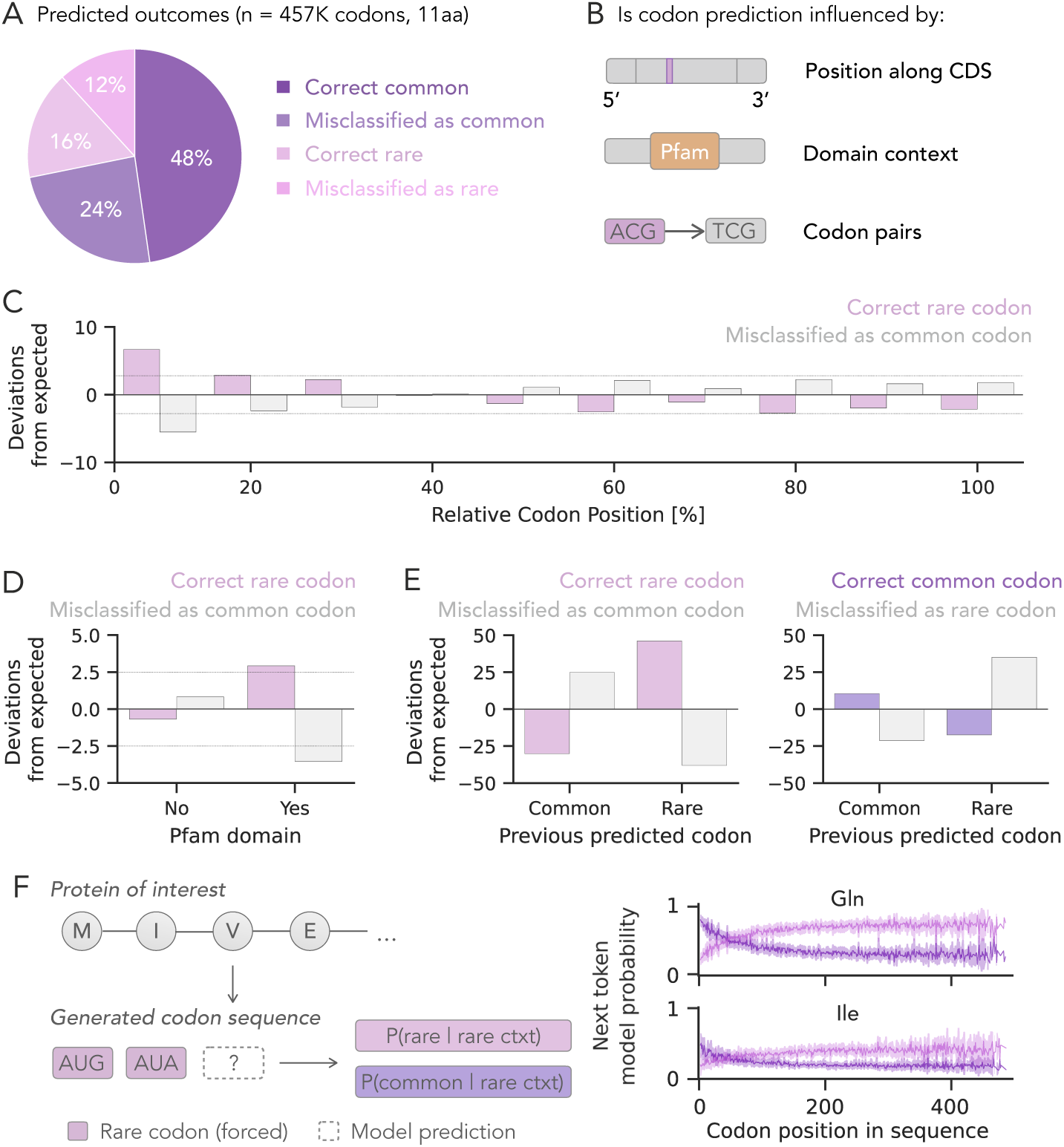
Contextual influences on codon prediction. (A) Codon predictions from the human test set were categorized into four classes based on whether the predicted codon was common or rare and whether it matched the wild-type codon. (B) To test whether codon predictions are influenced by sequence position, domain context, or previous codon identity (rare/common), we performed separate chi-squared tests for true common codons (correctly predicted common and misclassified as rare) and true rare codons (correctly predicted rare and misclassified as common). For each analysis, observed prediction frequencies were compared to expected frequencies under the assumption of independence between prediction outcome and the feature. (C–E) Standardized residuals are plotted to show deviations from expected frequencies. Dashed lines indicate residuals exceeding the significance threshold. Only significant feature-outcome analyses are shown. (C) Correct rare codon predictions were enriched near the 5’ end (first 10%), whereas misclassification of rare codons as common was reduced in this region. (D) Correct rare codon predictions were enriched within Pfam domains. (E) Rare codons were enriched after rare codons, and common codons were enriched after common codons. (F) Rare codon context experiment. Preceding codons were constrained to rare codons and the model probability of rare versus preferred codons was evaluated at the next position. Longer rare codon contexts increased the probability of continued rare codon prediction.

We first asked if the position in the overall coding sequence influenced predictions; rare codons are enriched towards the beginning of coding regions in eukaryotic genes, whether as a result of selection on translation or as a byproduct of other constraints (Tuller *et al*., 2010; Sejour *et al*., 2024). Trias learns to predict rare codons near the beginning of genes, with more correct predictions of rare codons as rare codons and fewer incorrect predictions of rare codons as common codons in the first 10% of the coding region (Figure 3C).

We also investigated whether codon predictions were affected by location within protein domains. The natural occurrence of rare versus common codons is similar within annotated Pfam domains and outside of domains. But, within domains, the model tends to correctly predict rare codons (Figure 3D). These results suggest that rare codon usage within domains is either more constrained or more learnable from distantly related sequences.

We then considered the impact of nearby sequence context. For both rare and common natural codons, the model’s prediction at a given position was associated with the prediction for the preceding codon (chi-squared test, p-value*<*1e-5). Rare predictions followed rare predictions more often than expected under independence, even where the natural codon was common, and common predictions followed common predictions more often than expected, even where the natural codon was rare (Figure 3E, Supplementary Figure S9).

Trias-generated sequences generally favor common codons, but a distinct subset of sequences are generated with rare codons, more closely resembling their wild-type counterparts (Figure 2A). Rare codons tend to cluster in natural sequences (Clarke and Clark, 2008) and we hypothesized that Trias may learn from such contexts. To test how rare codon context influences predictions, we designed an *in silico* experiment on the human test set sequences (Figure 3F): at each position, we set all preceding codons to the rarest synonymous codon (lowest sRSCU per amino acid) and compared the model’s probability of predicting the rarest versus most common (highest sRSCU) synonymous codon next. The preference for rare codons increased with the number of preceding rare codons (Figure 3F; Supplementary Figure S10). Thus, although Trias generally favors common codons, stretches of rare codons can shift the model toward continued rare codon use.

### Trias scores correlate with experimental measures of protein output

To evaluate the ability of Trias to predict protein output, we began by benchmarking its zero-shot predictions against published experimental data. A study from Bicknell *et al*. (2024) measured the protein output and half-life of 30 distinct mRNA variants encoding GFP, differing only in synonymous codon choice and spanning a range of codon optimality and propensity to form secondary structure (Supplementary Figure S11). These variants had been transfected into HEK293 cells as mRNAs with N1-methylpseudouridine (m1Ψ) replacing uridine, a standard base modification in therapeutic mRNAs. We scored each sequence with Trias by computing its length-normalized likelihood under the model, yielding a Trias score bounded between 0 (worst) and 1 (best). The Trias scores correlated strongly with the measured GFP fluorescence (Pearson’s *r* = 0.8) and mRNA half-life (Pearson’s *r* = 0.91) of the reporters (Figure 4A,B). Thus, despite being trained only on natural sequences and without being provided any information on mRNA translation or half-life, Trias learned codon patterns that align with the measured protein output and stability of synthetic reporters.

**Figure 4:**
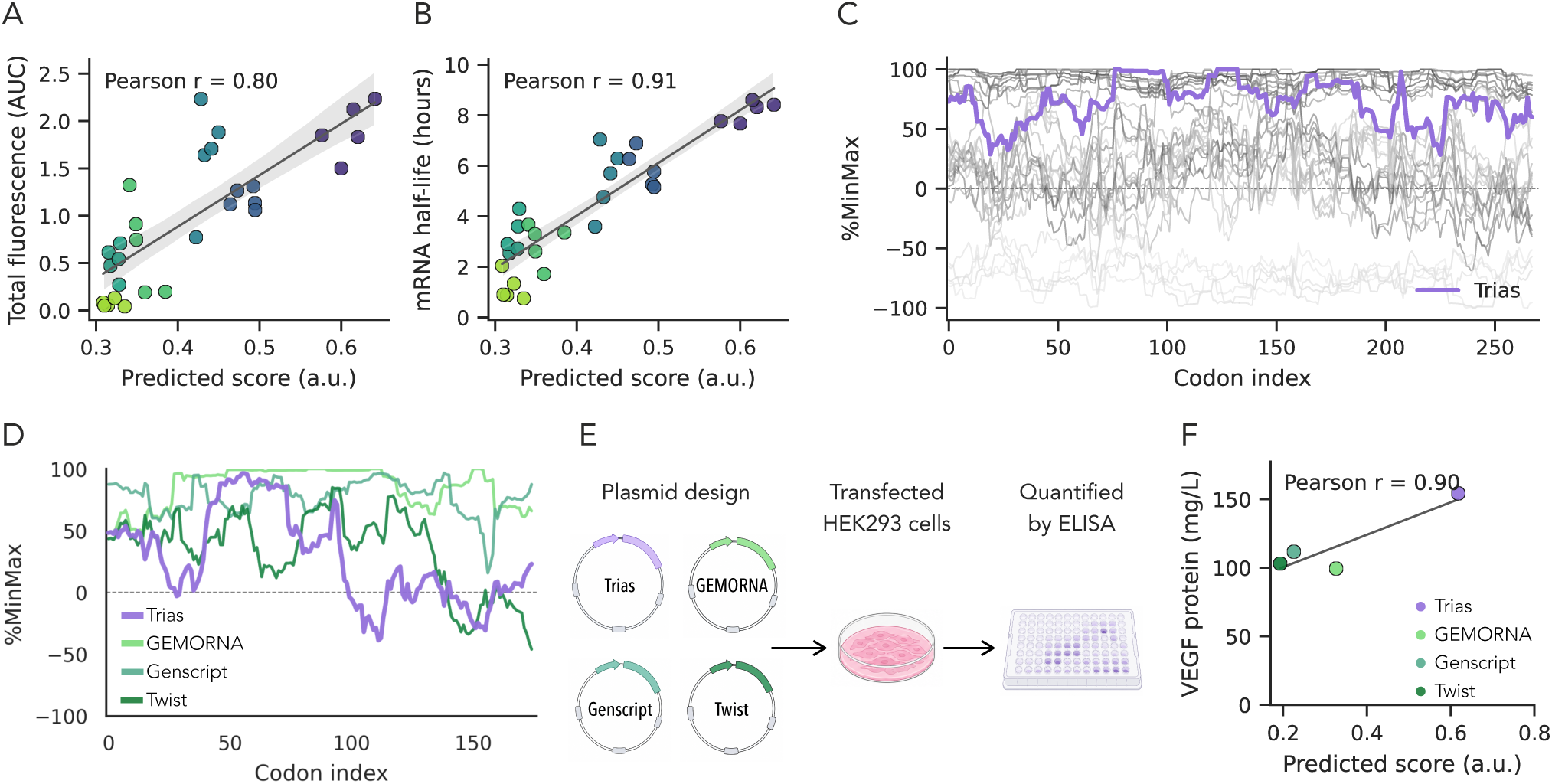
Trias-predicted codon sequences correlate with experimental data and differ from commercial codon optimization tools. (A,B) Correlation between Trias-predicted sequence likelihood and (A) total GFP fluorescence (AUC, area under the curve over time), and (B) mRNA half-life, for 30 GFP codon variants expressed in HEK293 cells (data from Bicknell *et al*. (2024)). (C) %MinMax codon usage profiles of all 30 GFP variants (gray) compared to Trias-generated sequence. (D) %MinMax codon usage profiles of VEGF sequences designed by Trias, GEMORNA, GenScript, and Twist. (E) Design and validation workflow: VEGF plasmids were transfected into HEK293 cells and quantified by ELISA. (F) Secreted VEGF versus Trias score across the four designs.

We then extended this zero-shot analysis to additional datasets measuring protein output of synonymous variants of several exogenous reporters in human cells and in mice (Leppek *et al*., 2022; Zhang *et al*., 2025; Bicknell *et al*., 2024). We assessed the correlation between the Trias score of each mRNA variant and its protein output in each sample. We repeated this evaluation with scores from our smaller, mammal-only model (Trias-small) and, to benchmark against existing methods, with scores from two other codon models, CodonTransformer (Fallahpour *et al*., 2025) and DeCodon (Naghipourfar *et al*., 2024). For the recently published GEMORNA method, whose scoring code is not publicly available, we used correlation values reported in the original study (Zhang *et al*., 2025). Finally, as a baseline, we assessed the correlations between protein output and simple mRNA properties: preferred codon usage, nucleotide composition, and structure propensity.

Our model Trias-small showed the highest average correlation across all datasets, tied with GEMORNA and followed closely by the full Trias model (Table 1). Our models performed substantially better than a simple metric of preferred codon usage, sRSCU, although quantification of mRNA properties outperformed deep learning models on some datasets (particularly those with m1Ψ modified mRNAs). These results demonstrate that Trias captures sequence features predictive of high protein output in a synthetic context from among the many functional pressures shaping natural sequences.

**Table 1:** Zero-shot performance across seven independent datasets. NL6h and NL24h: NanoLuc expression at 6h and 24h in HEK293 (Leppek *et al*., 2022). Fluc24h and Fluc48h: Firefly luciferase expression at 24h and 48h in HEK293T (Zhang *et al*., 2025); Fluc-AUC: Firefly luciferase AUC in mouse; GFP-AUC: GFP fluorescence AUC in HEK293; mRNA-HL: GFP mRNA half-life in HEK293 (Bicknell *et al*., 2024). Best performance per dataset is shown in **blue**, second best in light blue.

|  |  | Unmodified mRNA |  |  | m<sup>1</sup>Ψ modified mRNA |  |  |  | Mean |
| --- | --- | --- | --- | --- | --- | --- | --- | --- | --- |
|  |  | NL6h<br>(n=24) | NL24h<br>(n=24) | Fluc-AUC<br>(n=5) | Fluc24h<br>(n=8) | Fluc48h<br>(n=8) | GFP-AUC<br>(n=30) | mRNA-HL<br>(n=30) |  |
| *Codon language models* |  |  |  |  |  |  |  |  |  |
|  | Trias | 0.48 | 0.49 | 0.95 | 0.79 | 0.85 | 0.80 | 0.91 | 0.75 |
|  | Trias-small | 0.47 | 0.45 | 0.98 | 0.82 | 0.88 | 0.79 | 0.90 | 0.76 |
|  | GEMORNA | 0.44 | 0.48 | 0.96 | 0.84 | 0.87 | 0.81 | 0.92 | 0.76 |
|  | CodonTransformer | 0.53 | 0.50 | 0.74 | 0.42 | 0.51 | 0.70 | 0.87 | 0.61 |
|  | DeCodon | 0.42 | 0.35 | 0.75 | 0.82 | 0.90 | 0.53 | 0.56 | 0.62 |
| *Sequence-derived features* |  |  |  |  |  |  |  |  |  |
|  | 3rd pos. GC% | 0.38 | 0.60 | 0.85 | 0.79 | 0.78 | 0.83 | 0.88 | 0.73 |
|  | U% | -0.45 | -0.33 | -0.91 | -0.86 | -0.82 | -0.83 | -0.88 | -0.73 |
|  | MFE | 0.24 | -0.32 | -0.81 | 0.11 | -0.01 | -0.83 | -0.83 | -0.35 |
|  | sRSCU | 0.21 | 0.01 | 0.52 | 0.53 | 0.56 | 0.60 | 0.76 | 0.46 |

Values denote the Pearson correlation between model score and experimental measurements.

### Trias generates sequences with high protein output

A major question motivating our work is whether a model trained on natural sequences, which are rarely under selection for maximal protein output, would nonetheless design new sequences that maximize output. Our results so far suggest that Trias can design sequences for high output. For instance, it generated a hypothetical coding sequence with a score of 0.77 for the GFP reporter used above, notably higher than the scores of all 30 measured variants (Figure 4C).

(By definition, its own generated sequence will have the highest possible score; the more interesting observation is the magnitude of the difference.) The strong correlation between Trias scores and protein output in our zero-shot analysis suggested that we could extrapolate to predict high output from our own designed sequences.

Evaluation of mRNA design often relies on expression of fluorescence proteins for their ease of measurement, but therapeutically relevant mRNAs may have different constraints than mRNAs that encode the small, easily folded GFP protein. To test the ability of Trias to generate therapeutically relevant sequences with high expression, we designed a sequence encoding the vascular endothelial growth factor VEGF-A. Delivery of VEGF mRNA has undergone clinical trials for use in treating heart failure (Collén *et al*., 2022). While endogenous VEGF is subject to complex translational repression in response to cellular conditions (Arcondeguy *et al*., 2013), a therapeutic mRNA variant would ideally be translated at high efficiency.

We generated VEGF-A codon sequences with Trias, GEMORNA (Zhang *et al*., 2025), and commercially available codon optimization tools from GenScript^1^ and Twist^2^. Trias scored these other sequences substantially lower than its own generated sequence; interestingly, its own sequence used less-preferred codons (Figure 4D). The sequences were encoded on plasmids and transfected into HEK293 cells, and secreted VEGF protein concentration was assayed by ELISA (Figure 4E). We found a correlation between the Trias scores of the four variants and their protein output (Pearson’s *r* = 0.90; Figure 4F). The Trias-designed sequence produced the highest protein output. These results show that the design capabilities of Trias extend beyond simple reporters to a therapeutically relevant protein.

## Discussion

Foundation models of nucleic acid sequence have become promising tools for synthetic mRNA sequence design, with broad applications in recombinant protein production and mRNA-based therapeutics. Our generative model, Trias, learns codon usage directly from natural sequences, capturing biologically meaningful patterns beyond simple frequency-based approaches. Across independent datasets of synonymous variants measured in human cells and in mice, its zero-shot scores correlate strongly with protein output and mRNA half-life, and sequences designed by Trias for a therapeutically relevant protein produced more protein than those from other models. Trias performs well as a small model with a phylogenetically narrow training set, supporting the value of models tailored to a specific biological scenario.

Many applications of nucleotide foundation models for sequence design make the implicit assumption that higher-likelihood sequences will correspond to higher protein expression, but natural mRNAs are not necessarily optimized for high expression. Despite this conceptual difference, the success of Trias validates the approach of learning from nature in designing synthetic mRNAs for higher-than-natural expression. While Trias generally designs sequences with better-than-natural codons, the exceptions to this pattern are intriguing. In certain cases, Trias generates sequences that match the suboptimal codon usage of their natural counterparts. This must indicate that there is some learnable pattern distinguishing the codon usage of these genes, whether a stronger constraint or simply one that is easier to learn from the training examples. Further work may elucidate whether the suboptimal codon patterns of these genes are important for proper expression and function in either a natural or a synthetic context.

Ultimately, a complete model for synthetic mRNA design must incorporate regulatory features beyond the codon sequence itself. Untranslated regions (UTRs) play key roles in mRNA stability and translation efficiency (Mayr, 2017; Leppek *et al*., 2018), and the UTR sequence can affect the impact of codon choice (Lyons *et al*., 2026). Understanding these interactions could further refine the model’s ability to design synthetic mRNA sequences; recent work has shown that incorporating UTRs can improve predictions of mRNA properties and protein expression (Li *et al*., 2025).

A model of synthetic mRNA must also consider the impact of chemical modifications such as N1-methylpseudouridine (m1Ψ) that are necessary to reduce innate immune reactions to therapeutic mRNAs (Metkar *et al*., 2024). This modification, not found in natural mRNAs, may change codon-anticodon interactions and affect translation speed (Monroe *et al*., 2024; Rozman *et al*., 2026). Notably, in our benchmarking, the percent of U bases correlated better with protein output than did any deep learning method for several datasets in which all U bases were replaced with m1Ψ. The deep learning models, including Trias, were trained on natural sequences that evolved alongside standard nucleotide chemistry, and modified mRNAs may add distinct functional constraints that cannot be captured from evolution. Finetuning Trias on chemically modified sequences will be important to account for these effects. Future work to expand the scope of our model can fully unlock the potential of generative models for mRNA design in synthetic biology and therapeutic applications.

## Methods

### Datasets

#### Training and test dataset

We downloaded all vertebrate sequences from the NCBI RefSeq database^3^ on August 5th, 2024. To ensure data quality, we applied the following filtering criteria: sequences labeled as “pseudo” were excluded, and only records containing the feature tags “CDS” and “translation” were kept. We verified that each sequence was complete, initiated with the start codon “ATG” and terminated with one of the three stop codons (“TAA” “TGA” or “TAG”). Additionally, the translated protein sequences were required to start with methionine “M” and end with a stop symbol “*”.

After processing the downloaded records, we filter the mRNA sequences according to the length of the coding region. This step resulted in approximately 26 million sequences. To reduce redundancy, we clustered sequences with 90% or greater nucleotide similarity using MMSeqs2 and kept only one representative sequence per cluster. This step reduced the data set to around 10.2 million sequences.

To evaluate model performance, we created a train-test split from our curated dataset of 10.2 million vertebrate RefSeq coding sequences. Our goal was to ensure that the test set of human genes did not have close homologs in the training set. To accomplish this, we set aside a test set and then removed close mammalian homologs from the training set, those with regions of at least 70% protein identity, a distance at which protein identity no longer implies codon identity in the absence of selection on synonymous sites (Warren *et al*., 2008).

The test set was selected from the human MANE annotation of one representative transcript isoform per gene. To ensure a diverse range of expression profiles, we randomly selected equal numbers of genes from each of three expression categories (high, medium, and low) based on GTEx expression levels. These transcripts were translated into protein sequences and used to query the full mammalian RefSeq dataset using BLAST (blastp -matrix BLOSUM80 -max_target_seqs 5000 -evalue 1e-5) to identify close mammalian homologs. We then identified hits with 70% protein identity within the aligned region and removed all mRNA isoforms of these genes from the training dataset. (As -max_target_seqs 5000 truncated the results for some test set queries, we removed these genes from the test set.) The final test set comprised 1078 human genes with coding sequences of less than the model’s 2,046 amino acid limit.

The final vertebrate training set consisted of all 10 million remaining sequences from the curated dataset after removing those with significant protein similarity to the test genes. To monitor training progress, we also randomly sampled 1% of the training data as a validation set, with sampling balanced across species.

We then created a mammal-only training set by selecting all 3 million mammalian sequences from the vertebrate training set. This mammalian training set is compatible with the same human test set.

#### Gene expression data from GTEx

The GTEx dataset provides gene expression data across 54 human tissues from approximately 1,000 individuals, measured via bulk RNA sequencing (RNA-seq). For this study, gene expression data were obtained from the GTEx Portal^4^ on December 11th 2024, from the GTEx_Analysis_2017-06-05_v8_RNASeQCv1.1.9_gene_median_tpm.gct.gz file, which contains median gene-level TPM values per tissue. Genes with no expression (TPM = 0) in any tissue were excluded. We then computed the median TPM across all tissues for genes in our human test set.

### Model architecture and training

Trias is based on the HuggingFace implementation of the BART encoder-decoder architecture (Lewis *et al*., 2019). The input sequence length is set to 2,048 tokens, with the first token reserved for a species tag to ensure species-specific optimization. The vocabulary consists of 64 codon tokens, 20 amino acid tokens, 640 species-specific tokens, and 4 special tokens (</S>, <UNK>, <PAD>, <MASK>).

The model uses the BartForConditionalGeneration class, with 15% of amino acid tokens in the protein sequence masked during training. Flash attention is applied to improve memory efficiency and computational speed. Optimization is performed using the Adam optimizer with a cosine learning rate schedule and warm-up steps. The learning rate starts at 1e-4 and a weight decay of 0.01 is applied. Dropout is set to 0.1. The model is trained for 650,000 steps, corresponding to approximately two epochs.

Training is carried out on an NVIDIA A100 GPU with 80 GB of memory. The batch size is set to 8, and gradient accumulation is used with an accumulation factor of 4, resulting in an effective batch size of 32. The architecture consists of 6 encoder layers and 6 decoder layers, each with 8 attention heads. The hidden size of the feedforward networks is 2,048 for both the encoder and decoder. This configuration results in a model with approximately 47M trainable parameters.

The Trias-small mammalian model consists of 6 encoder layers and 6 decoder layers, each with 4 attention heads. The hidden size of the feedforward networks is 1024 for both the encoder and decoder. This configuration results in a model with approximately 12.3 million trainable parameters. The model was trained for 162,500 optimizer steps (6.7 epochs over 3,118,098 protein-CDS pairs) using AdamW with a peak learning rate of 1e-4 under a cosine decay schedule with a 1,948-step (1%) linear warmup, a weight decay of 0.01, and an effective batch size of 128 sequences at a maximum sequence length of 2048 with mixed-precision (fp16) training. Training took approximately 160 hours on a single NVIDIA A100 80GB GPU.

### Data analysis and performance metrics

#### Gene ontology enrichment analysis

To investigate functional constraints on codon usage, we performed a gene ontology (GO) enrichment analysis on genes with predicted low and high sRSCU values. We classified the human test genes into two groups based on their predicted sRSCU: genes with low sRSCU (*≤*0.86) and high sRSCU (>0.86). We then performed GO analysis to identify functional terms overrepresented in these subsets compared to the full set of 1078 human test genes. GO term enrichment was assessed using the gprofiler Python package, with the background set defined as all 1078 genes in the test set. We applied Bonferroni correction for multiple hypothesis testing and set a significance threshold of p-value < 0.05. GO terms related to molecular function, biological process, and cellular component categories were considered in the analysis.

#### Codon usage calculation

To quantify codon usage bias, we computed the scaled relative synonymous codon usage (sRSCU) for every codon in each species based on species-specific codon frequency in the training set. sRSCU normalizes codon frequency within synonymous codon categories, allowing direct comparisons across different amino acids. It measures how frequently a given codon is used relative to the expected uniform distribution of synonymous codons for an amino acid. The relative synonymous codon usage (RSCU) for a given codon *c* is calculated following Sharp *et al*. (1986):

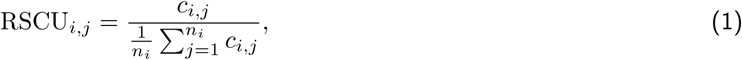

where *c_i,j_*is the observed frequency of the *jth* codon for the *ith* amino acid in the training set for a specific species, and *n_i_* is the number of synonymous codons for the *ith* amino acid. To derive sRSCU, we scale RSCU values by dividing by the highest RSCU value for each amino acid within a species:

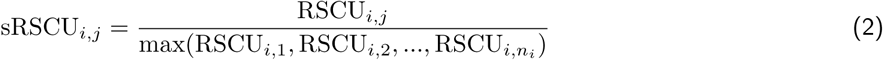

This ensures that the most frequently used synonymous codon receives a score of 1, while other synonymous codons are scaled accordingly. To calculate the sRSCU for a gene, we computed the average sRSCU value of all codons within that gene.

#### Expected value calculation for chi-squared tests

We used chi-squared tests to assess whether prediction outcomes were associated with specific sequence features (previous codon, codon position, and presence within vs outside a protein domain). Codon predictions from the test set were categorized into four classes (correctly predicted common, misclassified as rare, correctly predicted rare, or misclassified as common). Observed frequencies for all four prediction categories across each sequence feature are reported in Supplementary Tables S1–S4. For the chi-squared tests, we constructed separate contingency tables for true common codons (correctly predicted common and misclassified as rare) and true rare codons (correctly predicted rare and misclassified as common). Expected counts were calculated under the null hypothesis that codon prediction categories are independent of the tested feature. Expected values *E_ij_* were computed as follows:

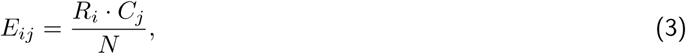

where *R_i_*is the total count for feature level *i*, *C_j_*is the total count for prediction category *j*, and *N* is the total number of codon predictions. To identify which feature levels contributed most to significant associations, we also computed standardized residuals.

#### Pfam domain annotation

We used InterProScan to identify protein domains in our test set. It scans protein sequences against a comprehensive collection of databases to annotate known domains, families, and functional sites. InterProScan was run locally on all human protein sequences in the test set. From the resulting annotations, we extracted Pfam domain matches and kept only those with e-values <1e-5 to ensure high-confidence predictions. The e-value indicates the number of matches expected by chance, with lower values reflecting higher significance. Codons were labeled as within a domain if they overlapped a detected Pfam domain, and as outside a domain otherwise.

#### Codon usage profiles

To analyze codon usage variability along sequences, we computed %MinMax profiles, which quantify deviations in codon usage relative to expected frequency distributions. This approach is adapted from Rodriguez *et al*. (2018) and provides a measure of codon bias by comparing observed codon frequencies within a sliding window (size = 18) to the expected frequency range. Positive %MinMax values indicate a preference for frequently used codons, while negative values reflect increased usage of rare codons.

To compare codon usage patterns between sequences, we used normalized Euclidean distance between their %MinMax profiles. The Euclidean distance was computed across the profile and normalized by sequence length. This approach provides a quantitative measure of similarity in codon usage fluctuations along the sequence.

#### Sequence likelihood scoring

To assess the optimality of a codon sequence *S* under our model, we define a scoring function *L*(*S*) as a length-normalized likelihood assigned by the model. Trias decodes codon sequences autoregressively, generating codons one at a time, where each codon is predicted based on the previously generated codons and the input protein sequence. The score of a sequence *S* = (*c*_1_*, c*_2_*, …, c_N_*) is given by:

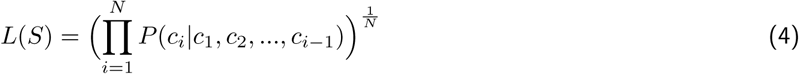

where *P* (*c_i_|c*_1_*, c*_2_*, …, c_i−_*_1_) is the model’s predicted probability for codon *c_i_*, given the preceding sequence at positions 1 through *i −* 1. In this score, the likelihood 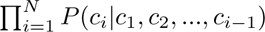 is normalized by sequence length *N* to create the geometric mean of the probabilities at each position. A *L*(*S*) value close to 0 indicates a sequence that is unlikely under the model, and an *L*(*S*) value close to 1 means that the model assigned high probabilities to codon choices at each position. A score of 1 corresponds to a sequence where the model is fully confident in every prediction (*P* (*c_i_*) = 1). Trias uses greedy decoding to generate codon sequences, a decoding strategy in which at each step the model selects the most probable codon. Greedy decoding ensures high-likelihood outputs, but not necessarily a global maximum.

#### Benchmarking

We evaluated each codon language model by scoring the benchmark coding sequences directly and correlating the resulting sequence scores with the reported experimental measurements using Pearson correlation. For Trias, CodonTransformer, and DeCodon, we used the geometric mean as model score computed from token-level logits. For GEMORNA, correlation values were taken from the original paper because the released code did not provide a comparable logits-based interface to score sequences. Reproducible scoring wrappers for CodonTransformer and DeCodon are provided in the lareaulab github repository.

#### Protein quantification

VEGF-A sequences were generated using Trias, GEMORNA, and codon optimization online tools from GenScript or Twist. Experimental validation was performed by SinoBiological. Each sequence was cloned into a pCMV3 expression plasmid and transfected into HEK293 cells using Sinofection reagent #STF02. VEGF-A secretion in the supernatant was measured by ELISA Sino (#SEK11066).

## Code availability

The code used to train Trias is available at https://github.com/lareaulab/Trias, and model weights are hosted on https://huggingface.co/lareaulab/Trias.

## Acknowledgments

This work was supported by grants from the National Institute of General Medical Sciences R01GM132104, the Rose Hills Foundation, the Shirl and Kay Curci Foundation, the Innovative Genomics Institute, and the Chan Zuckerberg Biohub (to L.F.L.). H.S. was supported by the Department of Defense through the National Defense Science & Engineering Graduate Fellowship (NDSEG) Program. We thank Kinsey Long for insightful input into the results.

## SUPPLEMENT

**Figure S1:**
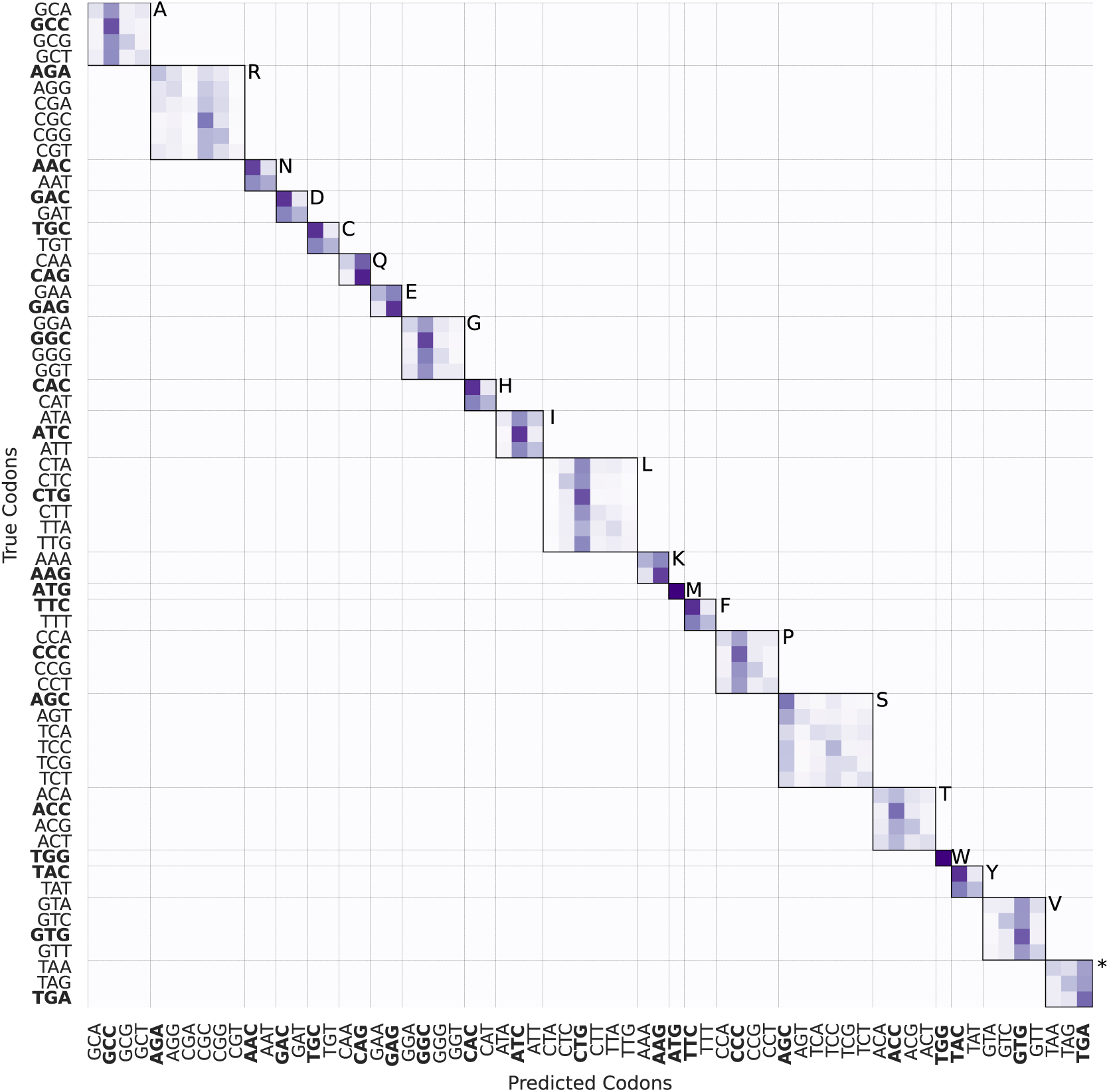
Percentage of Trias-predicted codons matching wild-type. Results are based on 683,080 codons from 1078 human test set sequences. Bold codons indicate the most frequent codon per amino acid, with synonymous codons grouped in boxes. The one-letter amino acid code is displayed.

**Figure S2:**
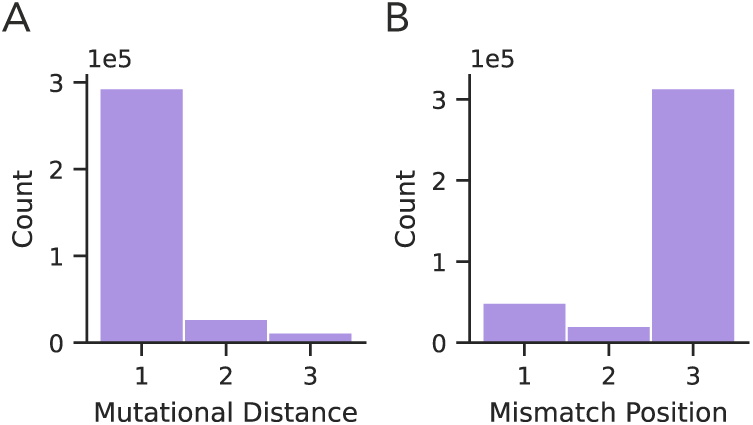
Codon mismatches and their positions. Among 683,080 codons in the human test set, approximately half differ from the wild-type codon, while only 0.001% result in an incorrect amino acid. (A) The majority of mismatches involve a single nucleotide substitution. (B) These substitutions predominantly occur at the third codon position.

**Figure S3:**
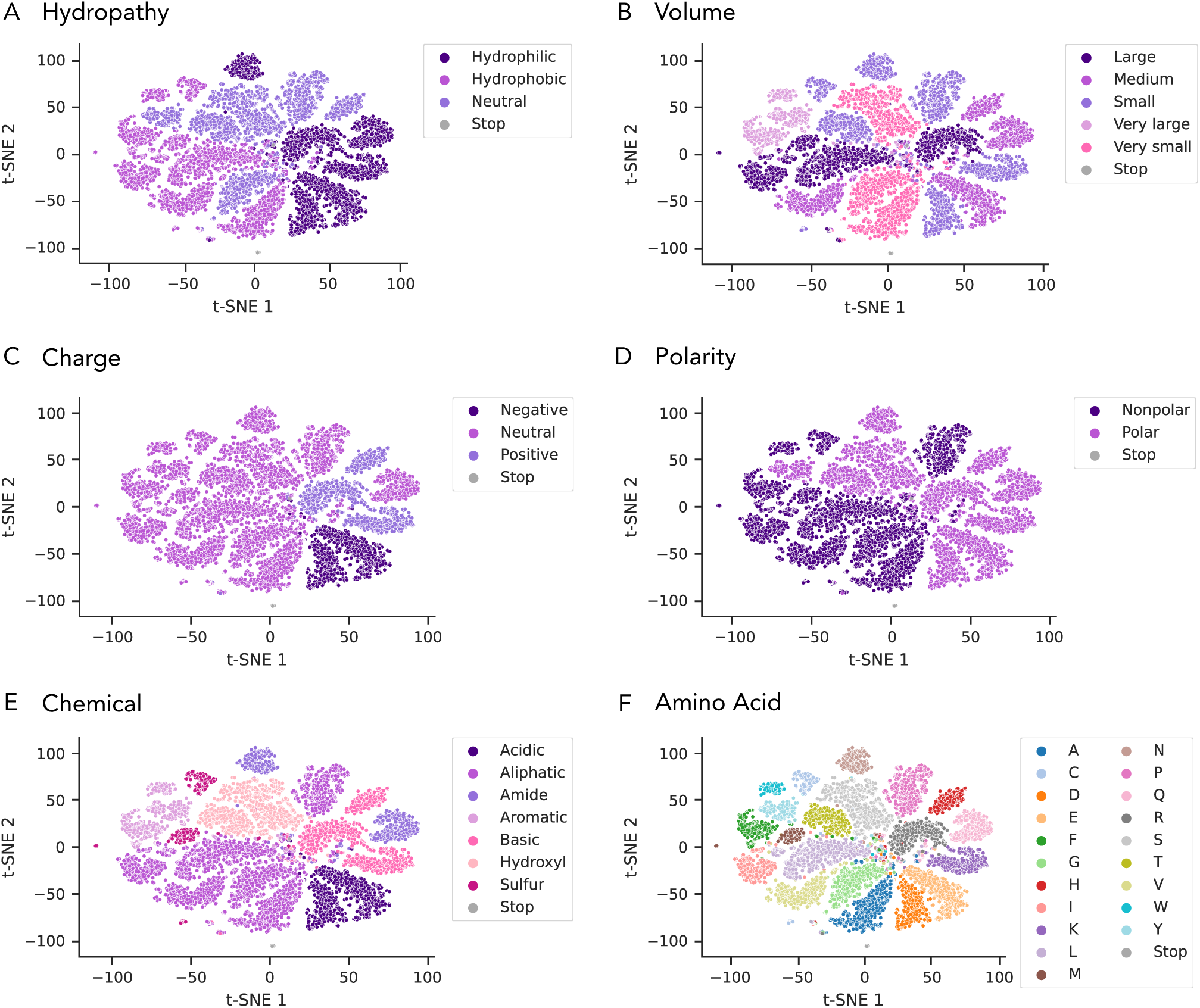
Projection of final encoder embeddings with different labels. Each dot represents an amino acid in 20 randomly selected protein sequences from the human test set. The same projection is shown with different labels based on classifications from this reference table^5^.

**Figure S4:**
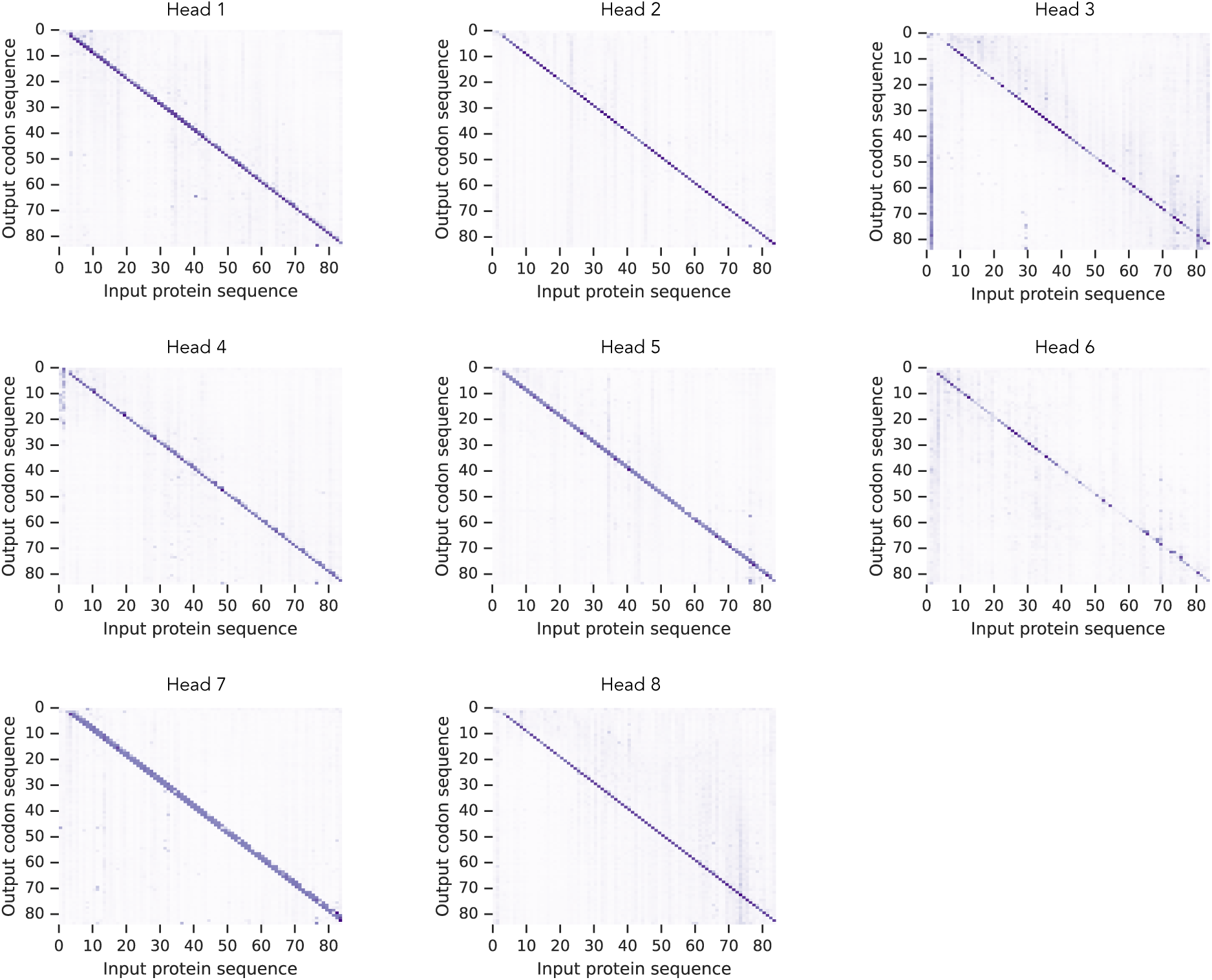
Cross-attention maps for all decoder heads averaged for COX7B2. Each panel shows attention from output codons (y-axis) to input amino acids (x-axis) for one decoder head. Attention weights are averaged across all decoder layers.

**Figure S5:**
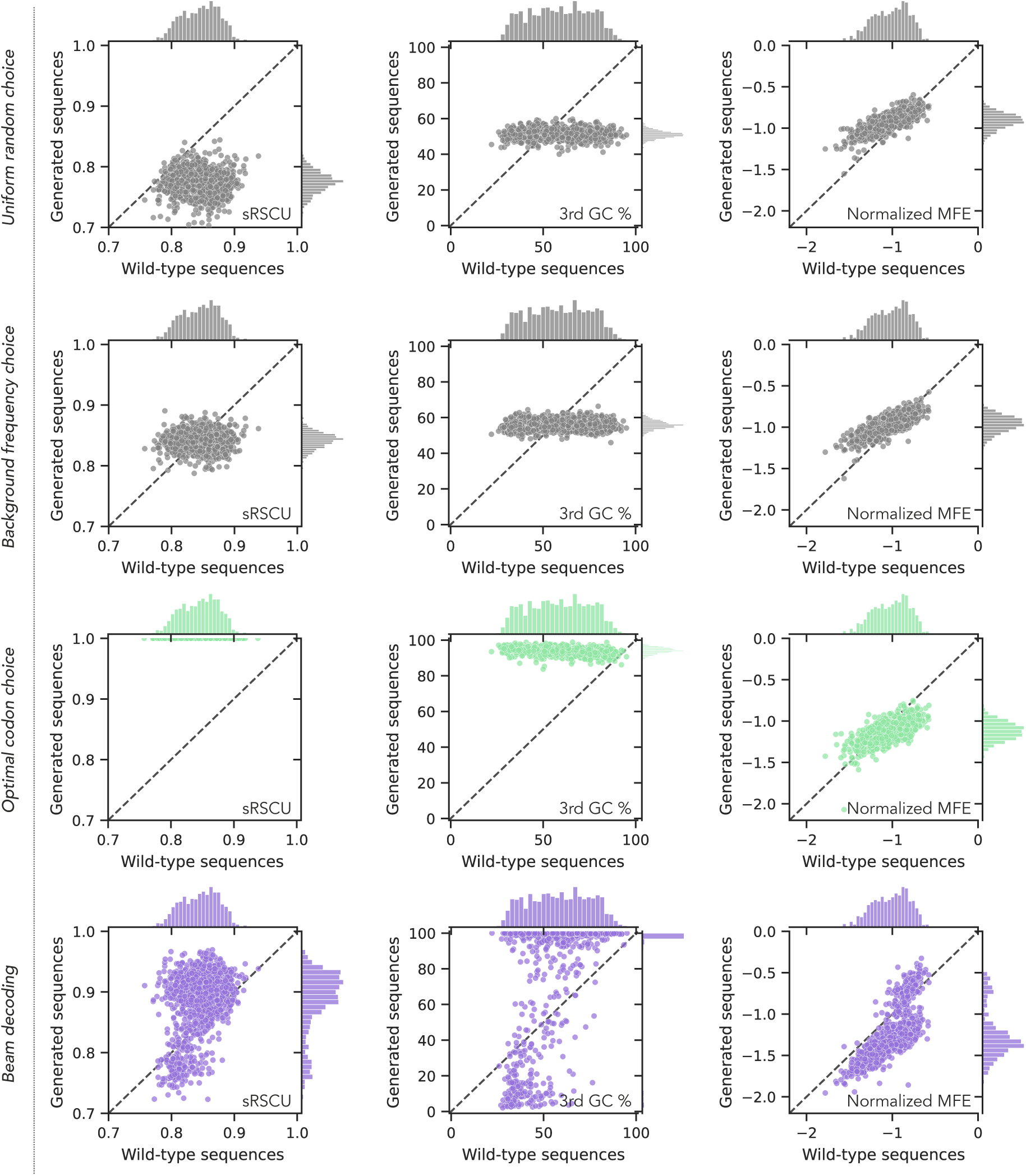
Comparison of generated sequences from random (uniform and codon-frequency–weighted sampling), baseline, and Trias using beam search decoding to wild-type sequences across scaled relative synonymous codon usage (sRSCU), GC content at the third codon position, and minimum free energy (MFE, normalized by sequence length) in the human test set (1078 sequences). The baseline selects the most frequent codon, while random models sample either from a uniform distribution or from background codon frequencies.

**Figure S6:**
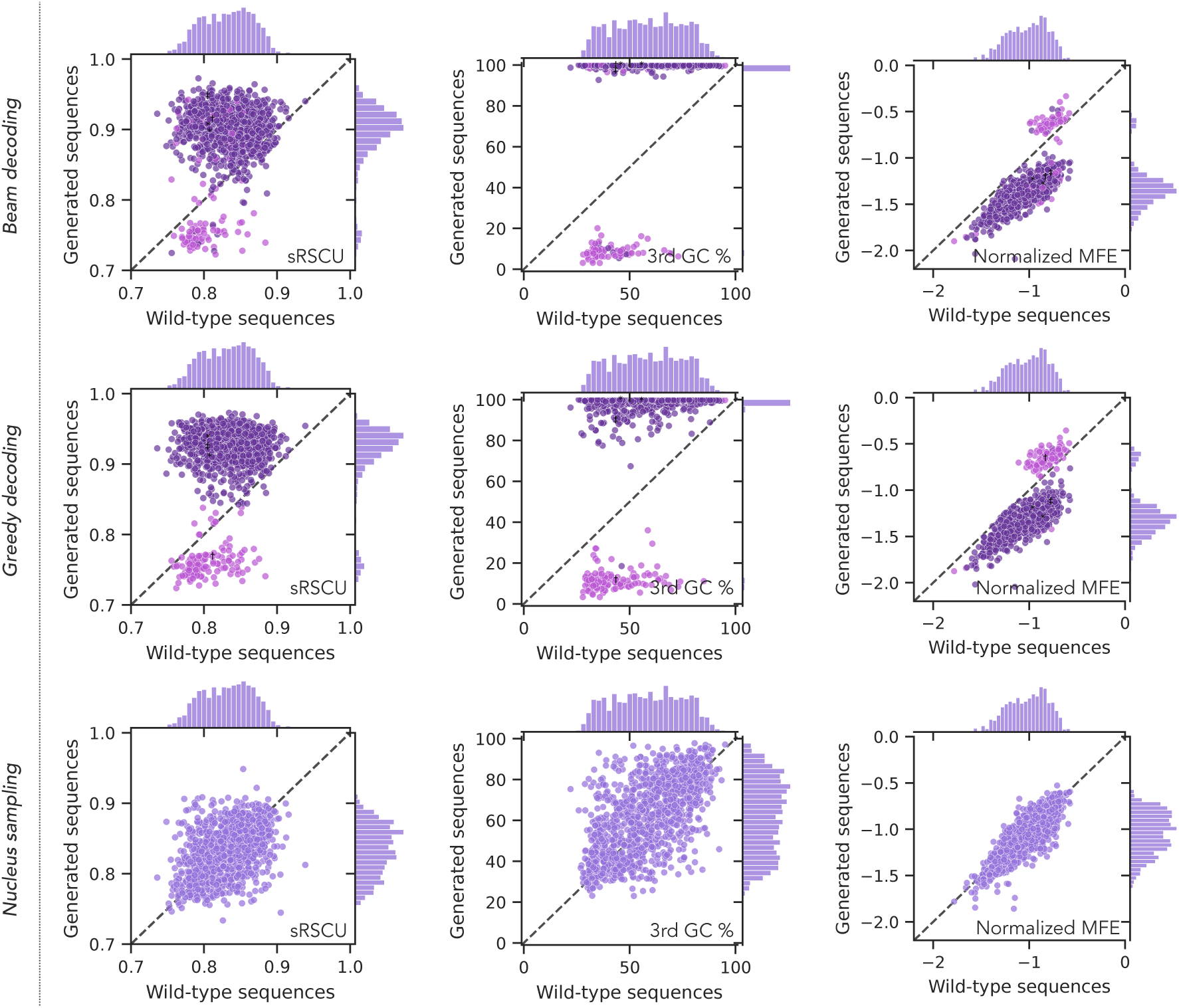
Sequence-level analysis of Trias-small-generated human codon sequences. Sequences from the mammal-only Trias-small model compared to their wild-type counterparts across sRSCU, third-position GC content, and normalized MFE, using beam search, greedy decoding, and nucleus sampling (human test set, 1,078 sequences).

**Figure S7:**
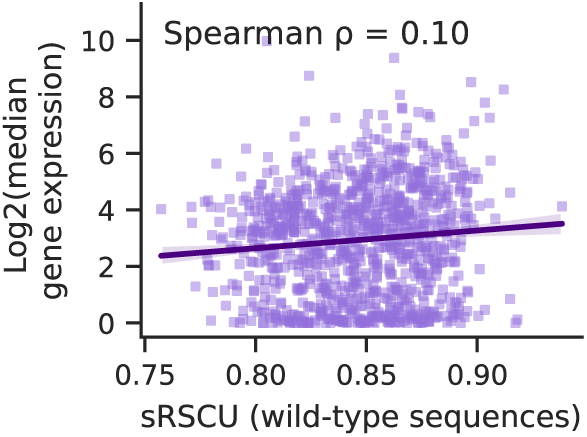
Log2-transformed median gene expression across all tissues in the GTEx dataset plotted against wild-type sRSCU values for 1020 human genes in our test set that matched GTEx entries.

**Figure S8:**
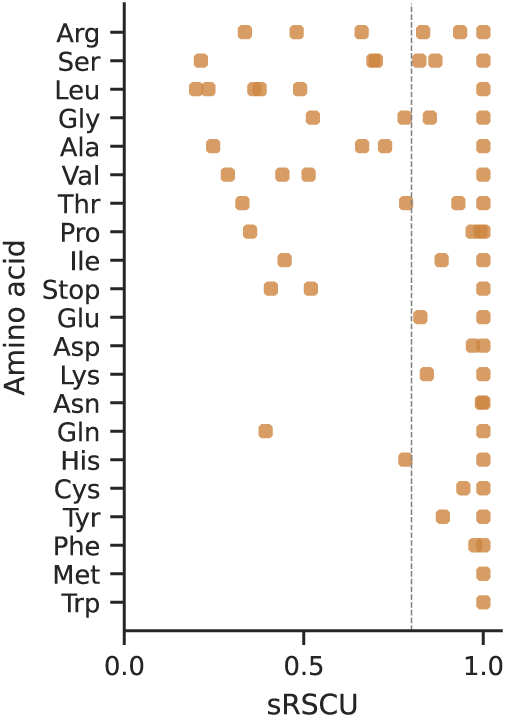
Corresponding sRSCU values for all synonymous codons across all amino acids. Rare codons were defined using an sRSCU threshold of <0.8

**Figure S9:**
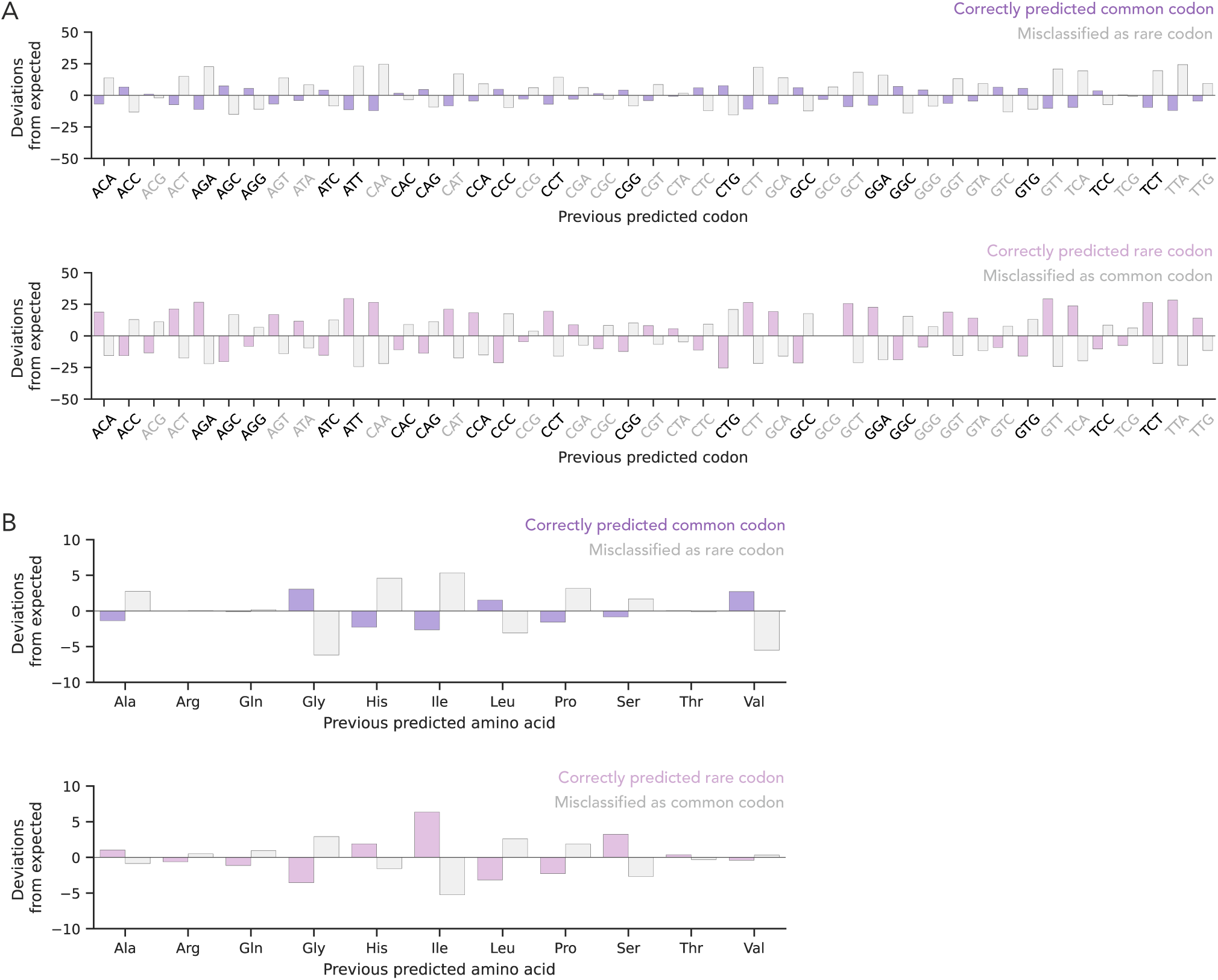
Influence of previous codon or amino acid on codon prediction. (A) Standardized residuals are plotted to show deviations from expected frequencies of next codon prediction classes based on the identity of the previously predicted codon. Rare codons are shown in gray, and common codons in black. (B) Same analysis as in (A), but grouped by the identity of the preceding amino acid.

**Figure S10:**
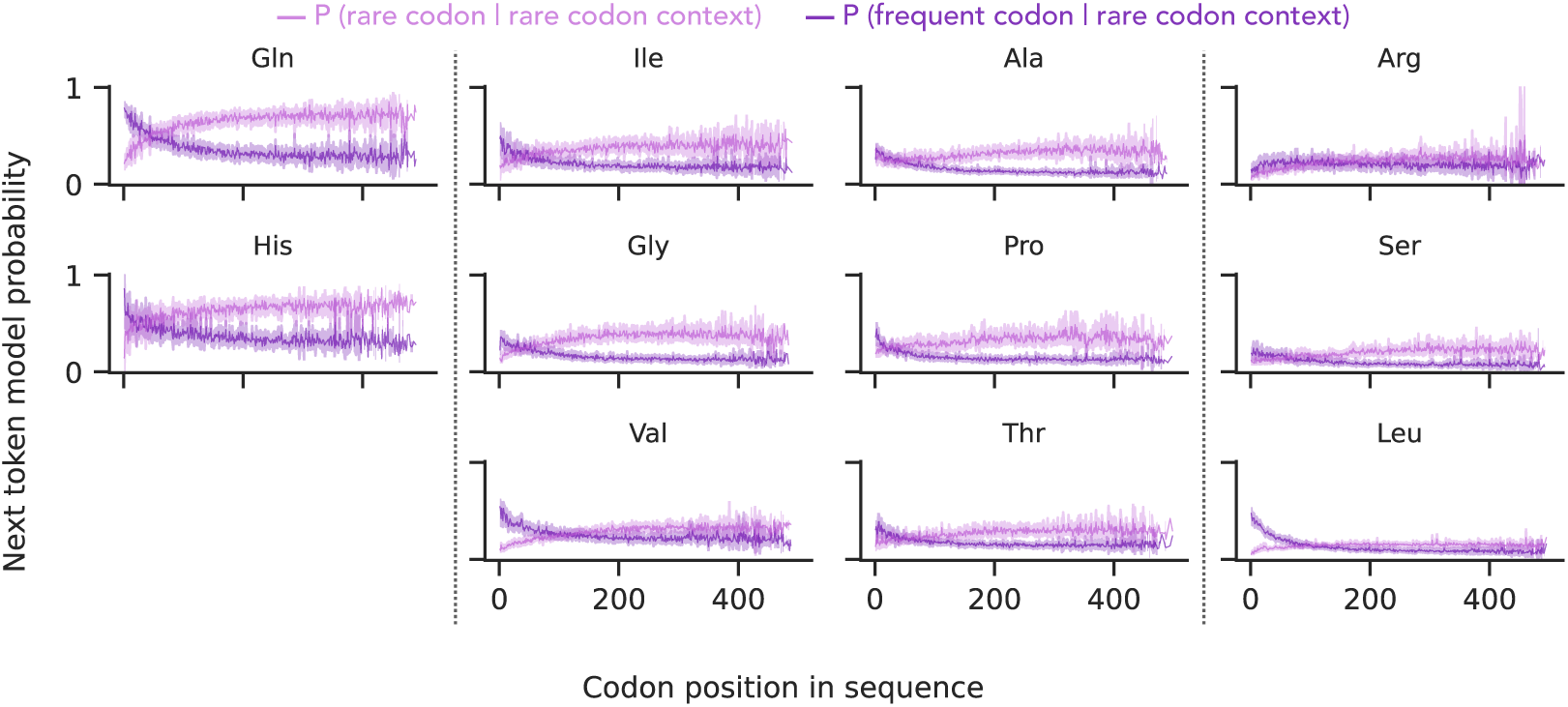
Rare codon context biases next token probabilities across amino acids. Model probability of selecting the next codon as frequent (highest sRSCU) or rare (lowest sRSCU), given that all preceding codons are rare. Amino acids are grouped by the number of synonymous codons (2, 3–4, and 6), with vertical dashed lines marking group boundaries. Mean and confidence interval are shown for 491 human test set sequences (sequence length *≤*500 codons).

**Figure S11:**
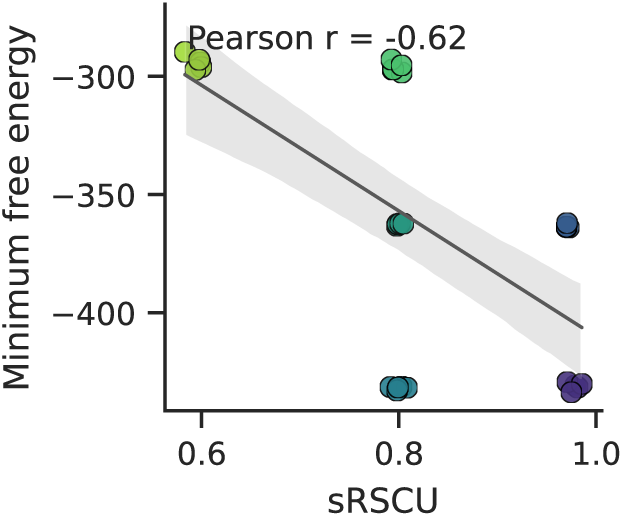
GFP codon variants from Bicknell *et al*. (2024). Minimum free energy (MFE) and codon optimality (sRSCU) values for 30 GFP variants. Variants were selected based on a grid of three distinct sRSCU and three different MFE values, with five variants chosen per region.

**Table S1:** Observed prediction counts by codon pair context. Counts of prediction categories for the second codon in each codon pair, grouped by whether the preceding predicted codon was rare or common.

| Prev. codon category | Correctly predicted common codon | Misclassified as rare codon | Correctly predicted rare codon | Misclassified as common codon |
| --- | --- | --- | --- | --- |
| Common | 163,528 | 35,122 | 45,530 | 83,934 |
| Rare | 54,603 | 18,781 | 29,370 | 26,041 |

**Table S2:** Observed prediction counts by amino acid. Counts of prediction categories grouped by the identity of the encoded amino acid. Only amino acids with both rare and common codons are included.

| Amino acid | Correctly predicted common codon | Misclassified as rare codon | Correctly predicted rare codon | Misclassified as common codon |
| --- | --- | --- | --- | --- |
| Ala | 22,509 | 5,822 | 7,854 | 11,302 |
| Arg | 18,636 | 4,611 | 6,237 | 9,282 |
| Gln | 16,132 | 4,002 | 5,575 | 8,399 |
| Gly | 21,445 | 4,743 | 6,670 | 10,527 |
| His | 7,942 | 2,220 | 3,009 | 4,163 |
| Ile | 12,984 | 3,591 | 5,380 | 6,794 |
| Leu | 33,357 | 7,893 | 10,812 | 16,703 |
| Pro | 21,337 | 5,562 | 6,825 | 10,495 |
| Ser | 27,549 | 6,984 | 9,991 | 13,880 |
| Thr | 16,991 | 4,188 | 5,938 | 8,644 |
| Val | 19,249 | 4,287 | 6,609 | 9,786 |

**Table S3:**
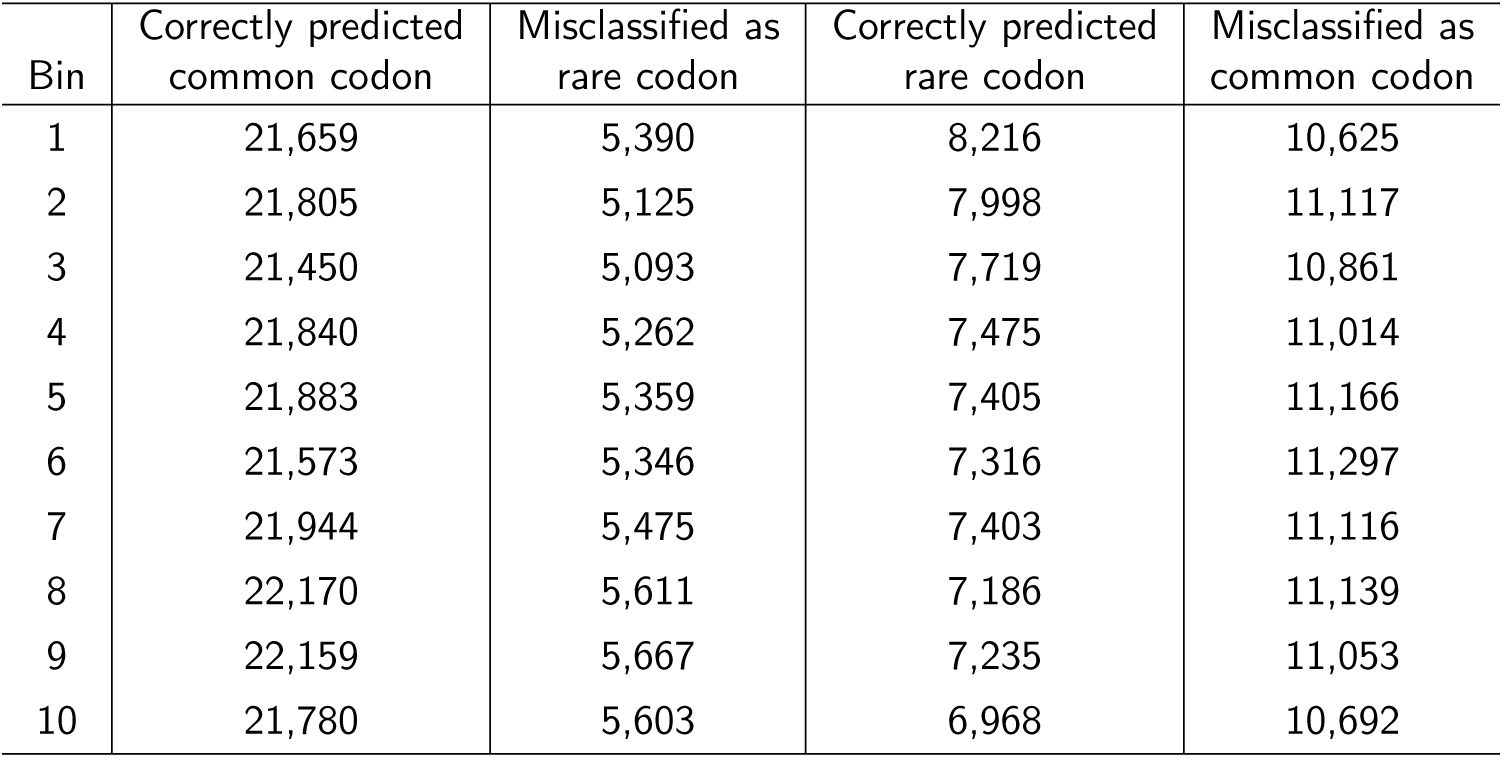
Observed prediction counts by relative sequence position. Counts of prediction categories across ten bins representing relative coding sequence positions (each bin corresponds to 10% of the total sequence length).

| Bin | Correctly predicted common codon | Misclassified as rare codon | Correctly predicted rare codon | Misclassified as common codon |
| --- | --- | --- | --- | --- |
| 1 | 21,659 | 5,390 | 8,216 | 10,625 |
| 2 | 21,805 | 5,125 | 7,998 | 11,117 |
| 3 | 21,450 | 5,093 | 7,719 | 10,861 |
| 4 | 21,840 | 5,262 | 7,475 | 11,014 |
| 5 | 21,883 | 5,359 | 7,405 | 11,166 |
| 6 | 21,573 | 5,346 | 7,316 | 11,297 |
| 7 | 21,944 | 5,475 | 7,403 | 11,116 |
| 8 | 22,170 | 5,611 | 7,186 | 11,139 |
| 9 | 22,159 | 5,667 | 7,235 | 11,053 |
| 10 | 21,780 | 5,603 | 6,968 | 10,692 |

**Table S4:** Observed prediction counts by domain localization. Counts of prediction categories grouped by whether the codon is located within or outside an annotated Pfam domain.

|  | Correctly predicted<br>common codon | Misclassified as<br>rare codon | Correctly predicted<br>rare codon | Misclassified as<br>common codon |
| --- | --- | --- | --- | --- |
| Pfam domain | 207,358 | 51,330 | 71,231 | 104,110 |
| Other | 10,905 | 2,601 | 3690 | 5,970 |

## Footnotes

1 https://www.genscript.com/gensmart-free-gene-codon-optimization.html

2 https://www.twistbioscience.com/resources/digital-tools/codon-optimization-tool

3 https://ftp.ncbi.nlm.nih.gov/refseq/release/

4 https://www.gtexportal.org/home/

5 https://www.imgt.org/IMflTeducation/Aide-memoire/_UK/aminoacids/IMflTclasses.html

